# Transcellular progression of infection threads in *Medicago truncatula* roots is controlled by locally confined cell wall modifications

**DOI:** 10.1101/2022.07.07.499094

**Authors:** Chao Su, Guofeng Zhang, Marta Rodriguez-Franco, Jenny Wietschorke, Pengbo Liang, Wei Yang, Leonard Uhler, Xia Li, Thomas Ott

## Abstract

The root nodule symbiosis with its global impact on nitrogen fertilization of soils is characterized by an intracellular colonization of legume roots by rhizo-bacteria. While the symbionts are initially taken up by morphologically adapted root hairs, rhizobia persistently progress within a membrane-confined infection thread through several root cortical and later nodular cell layers. Throughout this transcellular passaging, rhizobia have to repeatedly pass host plasma membranes and cell walls. Here, we genetically dissected this essential process and describe the concerted action of the symbiosis-specific pectin methyl esterase SyPME and the pectate lyase NPL at the infection thread and transcellular passage sites. Their coordinated function mediates spatially confined pectin alterations in the cell-cell interface that result in the establishment of an apoplastic compartment where bacteria are temporarily released into and taken up from the subjacent cell. This process allows successful intracellular progression of infection threads through the entire root cortical tissue.

## Introduction

Legumes evolutionary maintained the intriguing ability to intracellularly accommodate symbiotic bacteria called ‘rhizobia’. This mutualistic interaction results in the development of the root nodule symbiosis (RNS) that is morphologically initiated by a re-orientation of root hair (RH) growth, a process named ‘root hair curling’, followed by bacterial capture within a newly formed structure called the ‘infection chamber’ (IC, Fournier *et al*., 2015). The entrapped rhizobia start dividing inside the IC and trigger invasive growth of a tunnel-like structure, the ‘infection thread’ (IT) (Gage, 2004; Fournier *et al*., 2008). ITs are guided by pre-formed cytosolic columns (pre-infection threads) that are rich in endoplasmatic reticulum and cytoskeleton components (Timmers *et al*., 1999; Fournier *et al*., 2008, towards the basal membrane of the cell. In the course of IT progression, an organogenesis program is executed in root cortical and pericycle cells that results in the development of a nodule primordium (Hirsch, 1992; Xiao *et al*., 2014). ITs will transcellularly grow towards this primordium by penetrating several root cortical cell layers and finally release bacteria to cells inside the nodule. These colonized nodule cells provide the differentiated rhizobia (bacteroids) with an environment that is low in free oxygen and thus enables fixation of atmospheric dinitrogen gas by the rhizobial nitrogenase complex (Udvardi & Poole, 2013).

It can be assumed that spatio-temporally confined cell wall (CW) remodeling is required to initiate and maintain IT growth, transcellular passage of ITs as well as bacterial release. Cell walls of dicotyledonous plants mainly consist of cellulose, hemicellulose, pectins (rhamnogalacturonans), lignin and structural glycoproteins (Wolf *et al*., 2012). The polysaccharide callose is deposited in a more locally and temporarily confined manner and the loss of the callose degrading enzyme MtBG2 leads to defects in nodulation (Gaudioso-Pedraza *et al*., 2018).

Pectins are essential components of the primary cell wall and the middle lamella as found in young plant tissues. These heteropolysaccharides are synthesized in the Golgi apparatus and transported as methyl-esterified pectins to the plasma membrane where they are secreted into the apoplast (Voragen *et al*., 2009). Consequently, the methyl-esterified form of pectins serves as an abundant cell wall component of most root cell types. Immunolabelling of cell wall components in legumes revealed that de-esterified/un-esterified pectins delineate ITs (Vandenbosch *et al*., 1989; Rae *et al*., 1992; Xie *et al*., 2012; Tsyganova *et al*., 2019) while the IT matrix may predominantly be comprised of hydroxyproline-rich glycoproteins (HRGPs, extensins; Vandenbosch *et al*., 1989; Sujkowska-Rybkowska & Borucki, 2014; Tsyganova *et al*., 2019). While methyl-esterified pectins form a rather soft and gel-like matrix, de-esterification of pectins by pectin-methylesterases (PMEs) enables complexation by calcium ions into a mostly more rigid form called ‘egg-box dimer’. As PMEs form rather large and functionally redundant gene families, PME function has been assessed by using genetically encoded PME inhibitors (PMEIs) that simultaneously block PMEs at the site of PMEI accumulation (Wormit & Usadel, 2018). De-esterified pectins can also be targeted for degradation by pectate lyases (Wormit & Usadel, 2018). During RNS, a NODULE PECTATE LYASE (NPL), which is secreted via the VAMP721 pathway, regulates the stiffness of the CW by mediating the degradation of de-esterified pectins (Xie *et al*., 2012; Gavrin *et al*., 2016). As a consequence, most rhizobial infections are arrested at the IC stage in root hairs of *npl* mutant plants (Xie *et al*., 2012; Liu *et al*., 2019), indicating the importance of cell wall modifications for the infection process.

Here, we asked how cell wall modifications and IT transcellular passage through multiple cell layers is coordinated. To address this question, we genetically deciphered cell wall dynamics and identified a symbiosis-related PME (SyPME) that functions upstream of NPL activity. The concerted action of both enzymes and their differential and locally confined presence can explain the transcellular passage of ITs during symbiotic interactions.

## RESULTS

### Analysis of cell wall structures surrounding ITs

To study cell wall structure and composition during rhizobial infections and transcellular IT passage, we ran a set of pilot immuno-labelling experiments on indeterminate *Medicago truncatula* (hereafter Medicago) nodules using a series of selected antibodies against different cell wall constituents (Table S1). To identify ITs, we first labelled IT matrix glycoproteins using the MAC265 antibody (Vandenbosch *et al*., 1989). As expected, nodular ITs were specifically labelled by this approach while the peripheral cell wall of nodule cortex cells did not show any fluorescent signal (Fig. S1A). This was different when targeting xyloglucan as the most abundant hemicellulose by LM25. Here, we found a ubiquitous labelling of the cell periphery of all nodule cells including ITs (Fig. S1B). In contrast, different arabinogalactan proteins (labelled by LM2, LM14, LM30) that have been reported to serve functions during plant-microbe interactions (Nguema-Ona *et al*., 2013) were found to specifically accumulate within the infection zone and around symbiosomes (Fig. S1C-E, H-J’’). Next, we addressed the presence of different pectins including rhamnogalacturonans I (RG-I), homogalacturonans (HG) and rhamnogalacturonan II (RG-II). The linear (1-4)-β-D-galactan (recognized by LM5), an epitope of RG-I, was barely detectable inside the Medicago nodule sections (Fig. S1F). This is consistent with previously published data, where this epitope was almost absent in nodule sections from Medicago (Tsyganova *et al*., 2019). By contrast, (1-5)-α-L-arabinosyl, an epitope of RG-I that is recognized by LM6, is present in most cell walls of cells within the infection zone of the nodule (Fig. S1G, K-L’) and predominantly accumulates in the periphery of colonized cells of the fixation zone (Fig. S1M-M’), while uninfected cells within this zone did not accumulate (1-5)-α-L-arabinosyl (Fig. S1M’-M’’). To differentiate between HG subtypes, we applied two antibodies, LM20 and LM19, recognizing methyl-esterified and un-esterified HGs, respectively. While esterified pectins were present in most cell walls (Fig. 1A), un-esterified pectins predominantly accumulated in epidermal cells, outer cortical cells, and around infection threads, while the cell wall of central uninfected and infected cells were devoid of this processed form of pectin (Fig. 1C). Furthermore, and as shown for nodular tissue, un-esterified (Fig. 1D) but not methyl-esterified pectins (Fig. 1B) accumulated around ITs in root cortical cells. We further noticed that labelling of un-esterified pectins frequently extended slightly from the ITs towards neighboring cells.

**Figure 1:**
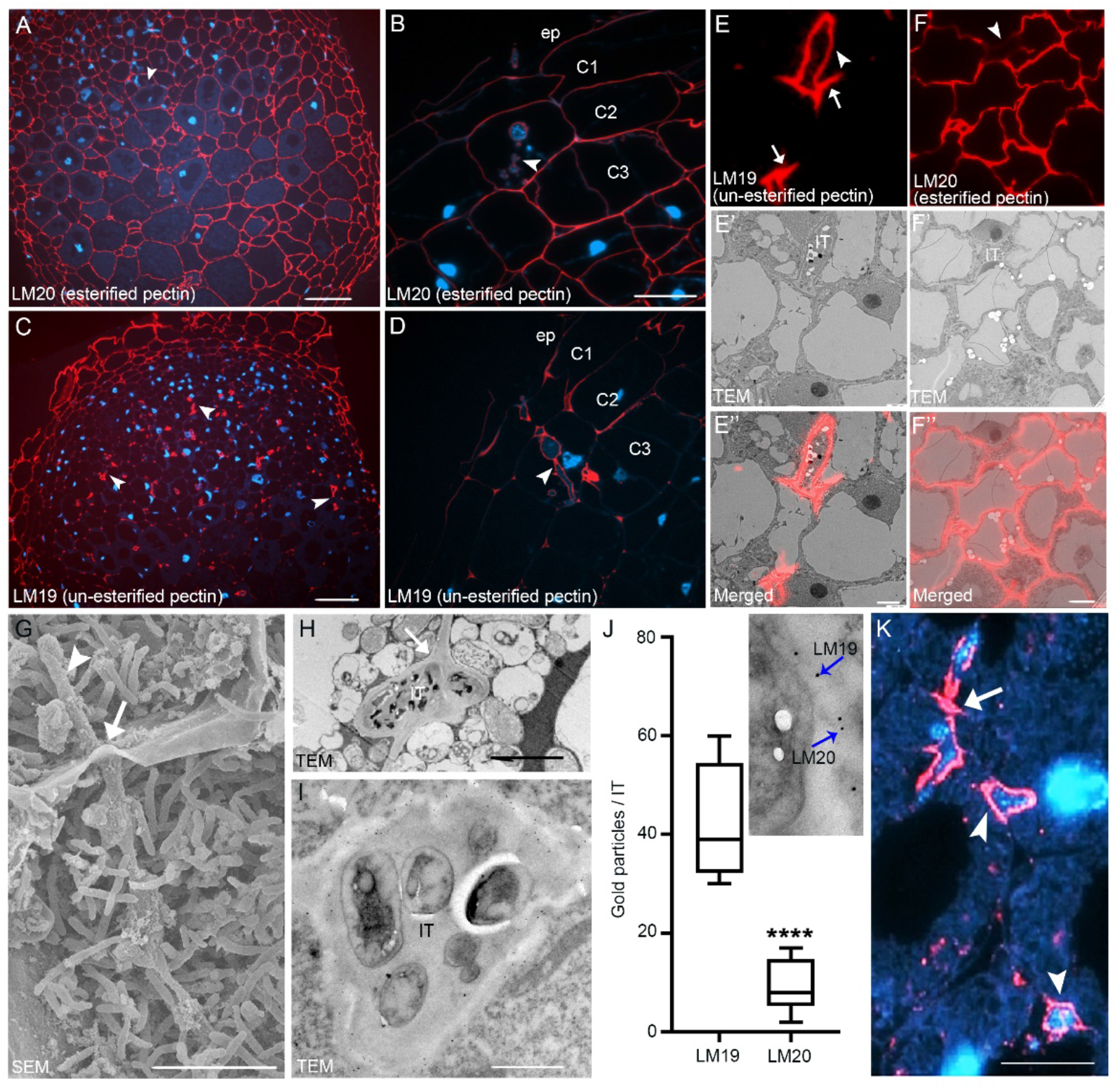
Infection-related modifications of pectins. (A-B) Esterified pectins (LM20, red) are present on the walls of all cell types in 14 days old Medicago nodules and root cortical cells but only weakly represented on the infection thread (arrowheads). (C-D) Enrichment of un-esterified pectins (LM19, red) in peripheral nodule cells and strong accumulation at nodular infection threads (C; arrowheads) and infection thread in root cortical cells (D; arrowheads). DNA was counterstained with DAPI (blue); ep, epidermis; C1, 1st cortical cell layer; C2, 2nd cortical cell layer; C3, 3rd cortical cell layer. (E-F’’) CLEM analysis of esterified (E-E’’) and un-esterified (F-F’’) pectins showing the fluorescence image after immunolabelling of ultrathin sections (E, F), the corresponding TEM image (E’, F’) and the corresponding merged images (E’’, F’’). Nodular infection threads (IT) are indicated by arrowheads (E, F) and extensions of pectin demethylesterification towards the direct neighboring cell at potential transcellular passage sites are labelled by an arrow (E). (G-H) Transcellular passage site (arrows) with a nodular infection thread (IT; arrowheads) passing from cell to cell as shown by scanning electron microscopy image (SEM) (G) and transmission electron microscopy (TEM) (H). (I) Double immune-gold labeling of un-esterified (LM19; 12 nm) and esterified (LM20; 5 nm) pectins at a nodular infection thread passage site showed labelling of the IT periphery. (J) Quantification data for double immune-gold labeling using LM19/LM20. Gold particles with different grain sizes were counted per IT (n=7). Data are means ± SE. Statistics were performed using an unpaired two-tailed t-test: ****p < 0.0001. (K) Strong enrichment of Ca^2+^-complexed pectins (labelled by the 2F4 antibody) around transcellular passage sites. DNA was counterstained with DAPI (blue). TEM: Transmission electron microscopy. SEM: scanning electron microscopy image Scale bars indicate 50 μm in A -D; 2.5 μm in E-F’’; 10 μm in G, H, I, and K.

### Assessing the ultrastructure of ITs using correlative light-electron microscopy

In order to dissect cell wall patterns at transcellular IT passage sites, we established a correlative light-electron microscopy (CLEM) protocol that allows searching for events by fluorescence microscopy (Fig. 1 E, F) and later to perfectly retrieve these sites in ultrathin sections using transmission electron microscopy (TEM) (Fig. 1E’, F’). Overlaying those images revealed that un-esterified pectins were present along the ITs and small segments of the host cell wall being in close proximity to the IT (Fig. 1E’’, F’’). These sites could be transcellular passage sites as frequently seen using scanning electron microscopy (SEM) (Fig. 1G) and TEM (Fig. 1H). Those observations were further confirmed by double immuno-gold labeling, which showed un-esterified pectins (LM19, 12 nm) being concentrated at the IT penetration site, while only a few gold particles were detected using LM20 (methyl-esterified pectins, 5 nm) (Fig. 1I, J). These images also revealed that CW structures at transcellular IT passage sites are fused and thickened (Fig. 1H). To assess whether this was a result of cell wall loosening and subsequent swelling or rather represents rigidified structures, we probed these samples using the 2F4 antibody, which recognizes ‘egg-box’ pectin dimers. Indeed, 2F4 immunofluorescence staining confirmed the accumulation of Ca^2+^ -complexed pectin around ITs and at the transcellular passage sites (Fig. 1K). This is in agreement with the observed enrichment of un-esterified pectins around ITs (Fig. 1C, D, E-E”), as Ca^2+^-complexation requires de-methylesterification of HGs.

### Identification of a symbiotic pectin methlyesterase

As pectin de-methylesterification is enzymatically mediated by pectin methyl esterases (PMEs; Pelloux *et al*., 2007; Jolie *et al*., 2010), we searched within the Medicago PME family, consisting of more than a hundred members, for suitable candidates. Using published transcriptome data (Breakspear *et al*., 2014; Damiani *et al*., 2016), we identified one gene (*Medtr4g087980* or *MtrunA17_Chr4g0069841*) to be consistently induced upon Nod Factor (NF) application and *S. meliloti* inoculation in roots (Fig. S2A). The gene also remained highest expressed in the nodule meristem (FI) and in cells of the distal infection zone (zIId) (Fig. S2B). Thus, we named it ‘*Symbiotic PME*’ (*SyPME*). We first verified the transcriptome data by *in situ* hybridizations and histochemical assay expressing a *GUS* reporter under the control of 2kb fragment upstream of the start codon as a putative promoter region. When hybridizing nodule sections with an antisense *in situ* probe, *SyPME* transcripts were found in cortical cells of nodule primordia (Fig. S2C) and in zone zIId of mature nodules (Fig. S2D). No signals were observed when using the sense probes as control in the same tissues (Fig. S2E, F). Accordingly, *SyPME* promoter activity, as delineated by β-Glucuronidase (GUS)-staining, was found within the entire cortex of nodule primordia (Fig. S2G), while a confined expression domain limited to the infection zone II was observed in young and mature nodules (Fig. S2H).

To assess the localization patterns of the SyPME protein, we initially generated a SyPME-GFP translational fusion, which was driven by the native 2 kb *SyPME* promoter fragment. Unfortunately, we were never able to obtain reliable fluorescent signals when using this construct in Medicago roots and nodules. However, when replacing the native promoter by a constitutively active Lotus *Ubiquitin 10* promoter, clear and confined fluorescence was observed in root hairs around the ICs (Fig. S3A) and along growing primary ITs (Fig. S3B). In line with this, nodular ITs were also decorated by the SyPME protein (Fig. 2A-F), while the strongest accumulations were observed at transcellular passage sites (Fig. 2D-F). Here, SyPME localization was strictly delineated to the peripheral cell wall at cellular conjunctions with crossing ITs (Fig. 2D-E), such sites that are rich in Ca^2+^-complexed un-esterified pectins (Fig. 1K). Furthermore, we noticed that SyPME also accumulated at both the tip region of growing ITs and a spatially confined site at the cell periphery that marks the site of the subsequent transcellular IT passage (Fig. 2A-C and Fig. S3E, G, G’).

**Figure 2:**
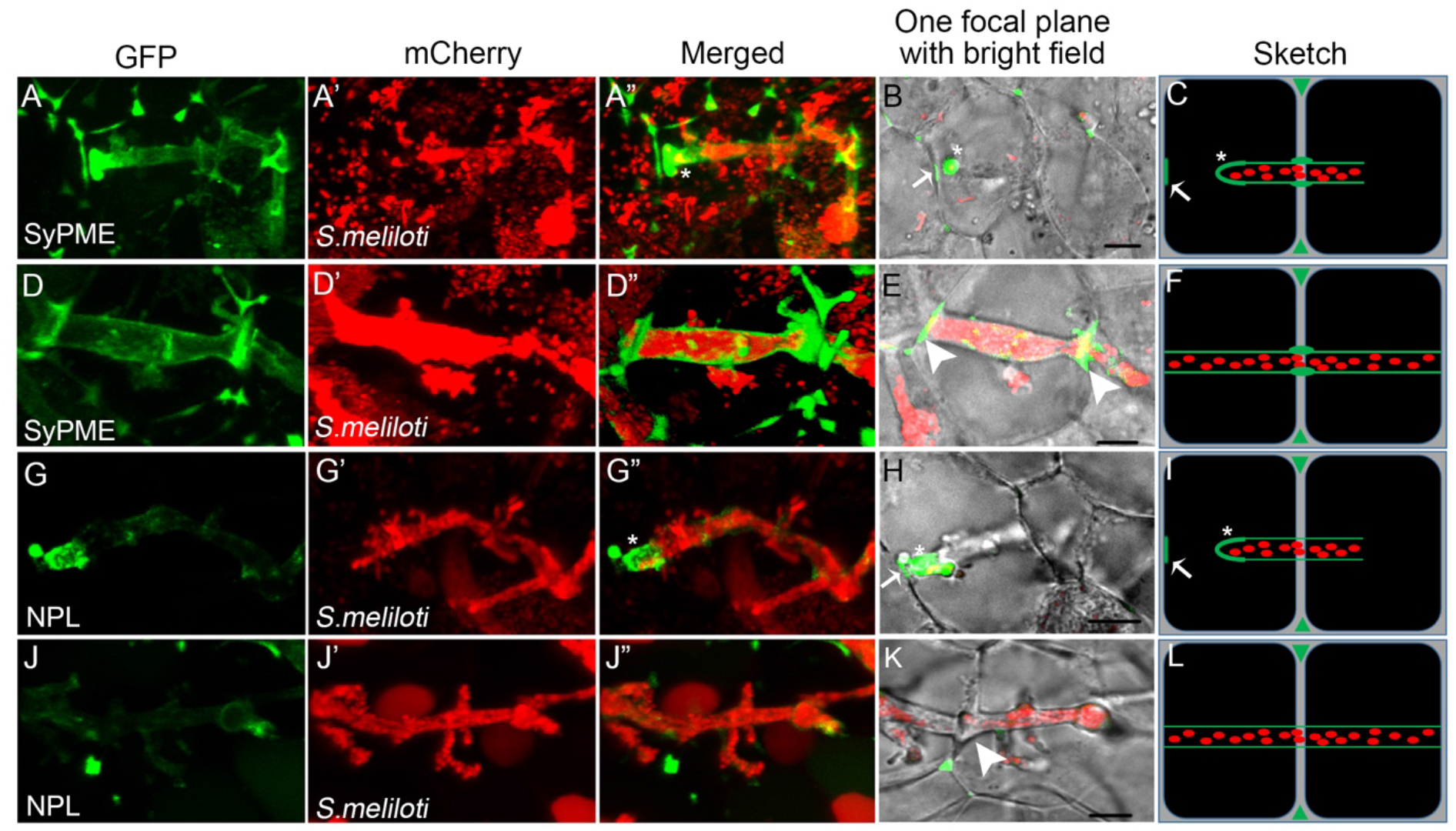
Spatially confined localization of the cell wall modifying enzymes SyPME and NPL. In 14 day old nodules obtained after inoculation with *S. meliloti* mCherry (red), SyPME-GFP (green) accumulates strongly at the tip and moderately at the shanks of nodular ITs (asterisk; A-C) and remains at transcellular passage sites beyond IT progression (arrowheads; D-F). In contrast, NPL-GFP (green) predominantly accumulates at the tip of nodular ITs (asterisk; G-I) but is absent from passage sites after IT progression (J-L). Both proteins also enrich at the passage site prior to IT arrival (arrow; B, C, H, I; for close-up see Fig. S3 G-H’). Scale bars indicate 5 μm. Images (A-A’’, D-D’’, G-G’’, and J-J’’) were processed as 3D projections using the Imaris software package.

### SyPME and NPL cooperate to regulate IT growth

As un-esterified pectins serve as substrates for pectate lyases (PLs) or polygalacturonases (PGs), we assessed the interplay between the Medicago NODULE PECTATE LYASE (NPL; Medtr3g086320 or MtrunA17_Chr3g0123331; Xie *et al*., 2012; Liu *et al*., 2019) and SyPME. Based on published data, *SyPME* and *NPL* are significantly co-expressed in infected root hairs (Correlation coefficient=0.9883, Fig. S2A). Compared to other members of the pectate lyase family being present in nodule transcriptomic data (Roux *et al*., 2014), *NPL* expression was found to be highest and mainly restricted to the nodule meristem (FI) and the zone zIId (Fig. S2B). This spatially controlled expression in nodules was also confirmed when generating a transcriptional GUS reporter using 2kb (2038 bp) upstream of the *NPL* start codon (Fig. S2I). On protein level and in line with patterns observed for SyPME, an NPL-GFP fusion protein driven by the endogenous *NPL* promoter was also localized to ICs, primary ITs in root hairs and the abovementioned spatially confined sites that will be penetrated by ITs (Fig. S3C, D, F). This suggests that SyPME precedes NPL-mediated pectin degradation at the IC, the tip region of growing ITs, and initially at the local cell wall site preparing for IT passage. Interestingly, older parts of these ITs showed reduced NPL accumulations (Fig. S3D) whereas SyPME protein levels remained high at these regions (Fig. S3B) suggesting a possible stiffening rather than loosening of the remnant ITs in root hairs. While NPL also localized to the tip of nodular ITs and local cell wall region near to nodular ITs (Fig. 2G-I and Fig. S3F, H, H’), the protein was absent from transcellular passage sites itself (Fig. 2J-L). Consequently, un-esterified pectins were, most likely, not degraded by NPL at these transcellular passage sites, confirming our hypothesis that these regions are stabilized and possibly sealed by Ca^2+^-complexed pectins provided by SyPME function. This would equally restrict apoplastic spreading of rhizobia as well as interference with other microbes colonizing the intercellular space of plants grown in natural soils.

### A genetic framework regulates IT growth and transcellular passage

To analyze the functional cross-talk between SyPME and NPL in more detail, we genetically assessed their impact on infection using loss- and gain-of-function approaches. In line with data reported for the *Lotus japonicus* (Lotus) *npl* mutant (Xie *et al*., 2012), significantly less nodules formed on the Medicago *npl* mutant compared to R108 WT plants (Fig. 3A). Furthermore, most infection events were aborted at the IC stage (Fig. 3B), an observation which is consistent with previously published data (Liu *et al*., 2019). To be able to differentiate between the requirement of NPL in the epidermis and the root cortex, we decided to additionally conduct an RNA interference (RNAi) approach and expressed the silencing construct under the control of different tissue-specific promoters in transgenic roots. We first expressed an empty control vector, where no changes in IT formation and progression were observed (EV; Fig. 3C, D). To generally prove the effectiveness of the system, we first used an NPL-RNAi construct driven by the Lotus *Ubiquitin 10* promoter. Constitutive over-expression of this silencing module frequently (37 out of 53) resulted in trapped rhizobia within the IC (Fig. 3E), while some ITs (10 out 53) manage to elongate but most likely stop in the root hairs (Fig. 3F). The same pattern was observed when driving the silencing construct by the *Solanum lycopersicum* (tomato) expansin 1 (*ProEXT1*) promoter that has previously been shown to mediate epidermis-specific expression in Medicago (Rival *et al*., 2012) (Fig. 3G, H). By contrast, cortex-specific silencing of *NPL* using the *Arabidopsis thaliana* endopeptidase *PEP*-promoter (*ProPEP*, Ron *et al*., 2014; Sevin-Pujol *et al*., 2017) predominantly resulted in normally developed ITs in infected root hairs that subsequently aborted in the root cortex (Fig. 3I, J). These results further demonstrate that NPL is required during both IT initiation and transcellular progression.

**Figure 3:**
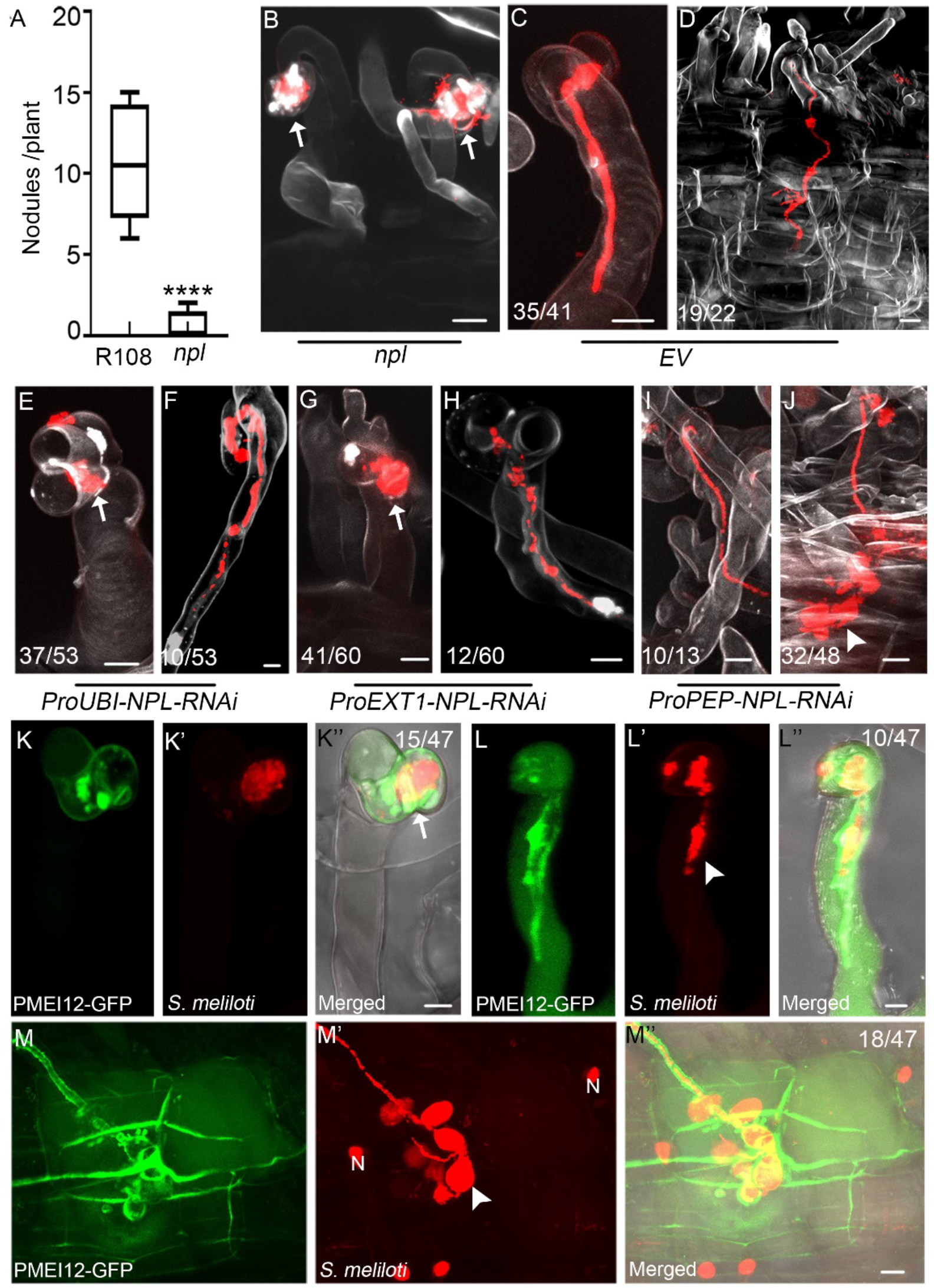
NPL activity is required throughout IT progression in different cell layers. The *npl* mutant developed significantly less nodules compared to the R108 WT control when grown for 7 days in the presence of *S. meliloti* in open pots (A; R108: n=12, *npl*: n=9) with the majority of infections already being blocked at the root hair stage (B). Data are means ± SE. Statistics were performed using an unpaired two-tailed t-test: ****p < 0.0001. WT-like IT growth in composite roots (C) and cortical cells (D) of control roots transformed with an empty vector (EV). Composite roots expressing an RNAi (RNA interference) construct against NPL (ProUBI-NPL-RNAi) showed aborted infections around the infection chamber (arrows; E) while some IT grew normally in root hairs (F). The same effect was observed upon epidermis-specific expression of the RNAi construct (ProEXT1-NPL-RNAi) (G, H). Silencing *NPL* expression in root cortical cells (ProPEP-NPL-RNAi) did not impair IT growth in root hairs (I) but transcellular progression in the root cortex (J). Expression of PMEI12-GFP under the control of the *NPL* promoter resulted in IT abortions around the infection chamber (K-K’’), in root hairs (L-L’’) and in the root cortex (M-M’’). N: nucleus; scale bars indicate 10 μm. All images are maximal projections.

To genetically test whether SyPME and/or other members of this protein family are required for successful infections, we first searched the Tnt1 transposon insertion collection and identified a single *sypme* allele carrying an insertion in the first intron (NF2281_high_35; Fig. S4A). Homozygous individuals, however, did not show any symbiotic phenotypes (Fig. S4B-F), which might be due to the large size of and functional redundancy within this gene family. In order to target functionally redundant PMEs with spatio-temporal precision, we expressed the *Arabidopsis thaliana PME INHIBITOR 12* (*PMEI12*) under control of the *NPL* promoter and inoculated these transgenic roots with *S. meliloti*. This protein has been previously demonstrated to efficiently inhibit PME activities (An *et al*., 2008; Lionetti *et al*., 2017). Indeed, the majority of infections were blocked at the IC stage (Fig. 3K-K’’) or during IT progression in root hairs (Fig. 3L-L’’) and the root cortex (Fig. 3M-M’’), thus phenocopying the *npl* mutant. This supports the proposed dependency of NPL mediated pectin degradation on preceding PME activity.

### Deciphering the steps of IT passage by CLEM

In a last set of experiments and to unambiguously unravel cell wall changes during transcellular IT passage, we used our CLEM approach to monitor pectin alterations within the cell-cell interface at different stages of transcellular IT progression. For this, we labeled un-esterified pectins (LM19, Fig. 4A) and the IT matrix (MAC265, Fig. 4B) and searched ultrathin sections (Fig. 4C-D) for ITs approaching the basal cell membrane/cell wall and the moment of IT passage. Un-esterified pectins accumulated around the future transcellular passage site, as defined by the cytoplasmatic column formed ahead of the IT, within the cell-cell interface (Fig. 4, stage I). This passage site further increased in size, expanded laterally and subsequently swelled in the central region (Fig. 4 stage II). This was followed by rhizobia entering a closed apopolastic compartment prior to entry into the neighboring cell that had already formed a pre-infection thread (Fig. 4, stage III). At the end, rhizobia entered the neighboring cell through the apopolastic space without any membrane confinement at this site (Fig. 4, stage IV).

**Figure 4:**
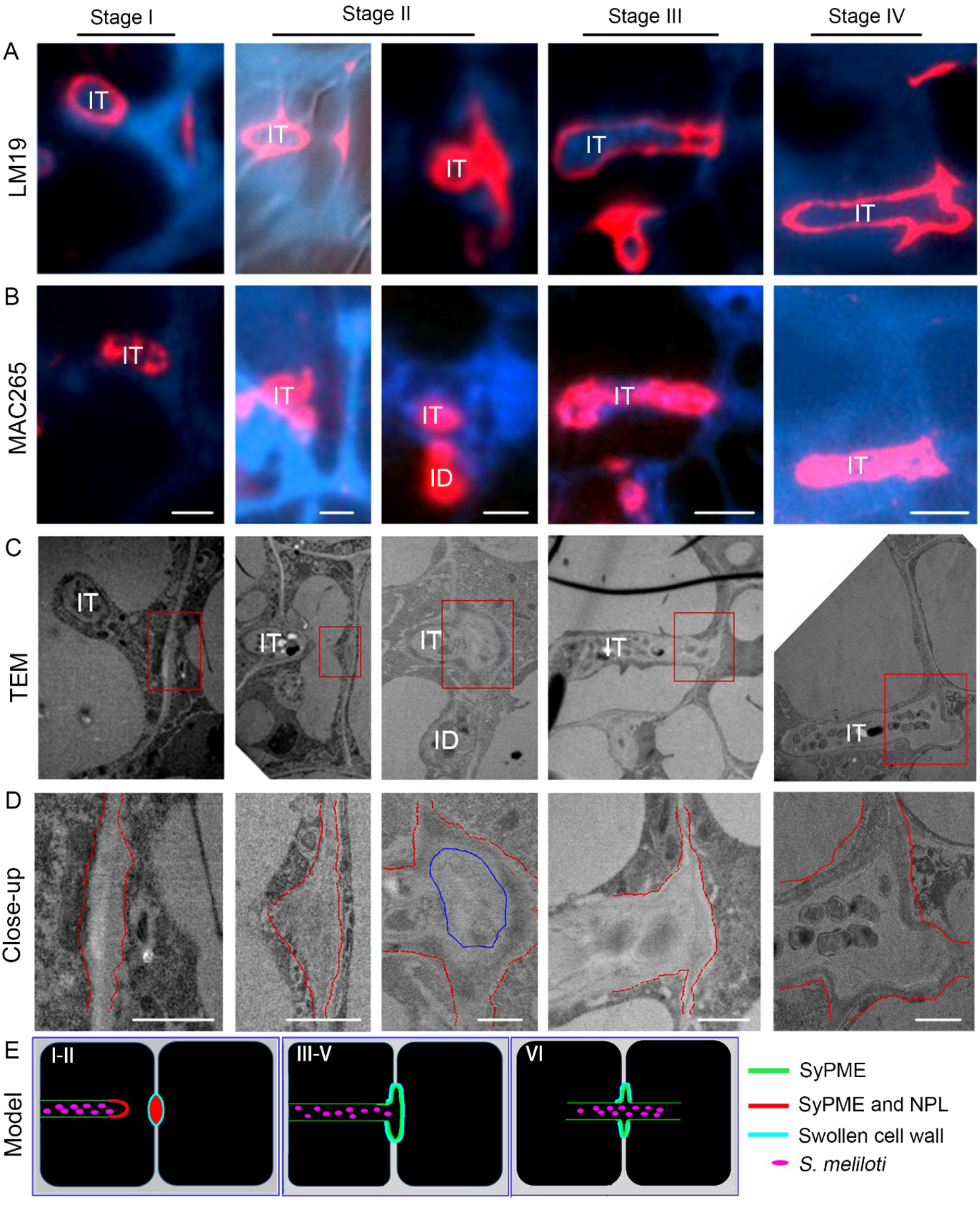
CLEM analysis of distinct steps during transcellular IT passage. CLEM analysis for cell wall modifications at the IT (infection thread) penetration sites for un-esterified pectins (labelled by LM19 (red), A) and IT matrix components (labelled by MAC265 (red); B) with consecutive sections. (C) Corresponding TEM micrographs and their close-ups (D) as indicated by the red boxes in (C). Red dashed lines (in D) indicate the border of cell wall; blue encircled region indicates the cell wall swelled in the central region. (E) Proposed graphical model for different stages of IT transcellular passage. Scale bars indicate 4 μm in (A-C) and 1 μm in D. ID: infection droplet.

Given the estimated diameter of the passage site in the range between 1 and 3 μm, the discontinuation of the IT membrane upon fusion of the IT with the basal membrane and the *de novo* invagination at the neighboring cell, we conclude that transcellular passage is a successive series of events (Fig. 4E). This includes 1) targeted secretion of PMEs and initially of NPL, 2) local pectin de-methylesterification and partial pectin degradation at the tip of the IT and at the local host cell wall prepared for penetration, 3) maintenance of SyPME but reduction of NPL protein levels at the passage site, 4) confined cell wall swelling at the passage site, 5) spatial release of rhizobia into a sealed apoplastic compartment and 6) uptake into the neighboring cell.

## Discussion

The intracellular colonization of differentiating nodule primordium cells is a conserved process in model legumes such as Lotus and Medicago as well as most agriculturally relevant legume species including soybean, pea and beans. This phenomenon requires the transcellular passage of the IT throughout several cell layers. While it has been assumed decades ago that ITs pass the cell wall via apertures around the middle lamella (McCoy & Russell, 1932), it has been proposed that this does not involve cell wall pits (plasmodesmata; Goodchild & Bergersen, 1966). This was further supported by the fact that the IT membrane fuses with the basal plasma membrane of the host cell upon transcellular passage (Szczyglowski *et al*., 1998). In indeterminate nodulators such as Medicago, transcellular progression of ITs does not only occur in the outer root cortex but also constantly in the distal part of the so-called nodular ‘infection zone’ (Gage, 2004). Here, we often observed a local thickening of the cell wall (Fig. 4C, stage I-II) at the future passage sites, while we never detected the formation of true cell wall apertures at these sites of IT penetration. Additionally, both SyPME and NPL accumulated at the prospective passage sites prior to IT arrival at the plasma membrane (Fig. 2B, H, and Fig. S3E-H’), which indicates that a spatially confined and partial pectin degradation is involved in preparing these sites for penetration by generating a weakened cell wall region. Subsequently, cell wall swelling at the penetration site (Fig. 4) might be induced by PME-mediated de-methylesterification of pectins, which can lead to cell wall expansion and hydration (Peaucelle *et al*., 2015). It should be noted that this process, alike the formation of egg-box pectin, also requires Ca^2+^-complexation (White *et al*., 2014; Wang *et al*., 2015). Consequently, the presence of complexed de-methylesterified pectins, as labelled by the LM19 and 2F4 antibodies (Fig. 1) around the transcellular IT passage sites, could result in local stiffening or swelling of the cell wall in the absence of NPL at these sites (Fig. 2J-K). Similar cell wall swellings were observed during colonization of roots by symbiotic arbuscular mycorrhiza fungi (Rich *et al*., 2014) and during stomata pore formation (Nadeau & Sack, 2002). While the initial processes might be comparable to those described here for rhizobial infections, the final stomatal pore is formed by the continuous accumulation of de-esterified pectins and polygalacturonase PGX3/1 activity (Rui *et al*., 2017). During transcellular IT passage, however, NPL only transiently accumulates prior to and upon the IT reaching the basal membrane, while PME accumulates throughout the process and remains present at this site (Fig. 2).

Swelling of the cell-cell interface may also be supported by a low cellular turgor pressure. While the cellular turgor needs to be reduced prior to cell divisions of root cortical cells during formation of the nodule primordium, it is likely to be still low at newly divided cells originating from the nodule meristem and it is only these newly divided cells that can be successfully entered by the IT (Monahan-Giovanelli *et al*., 2006). This process of cell wall loosening and swelling is, most likely, further mediated by expansin proteins. These apoplastic proteins support cell wall loosening without having any known enzymatic activity (Cosgrove, 2016). Indeed, apoplastic accumulation of expansins around ITs has been reported in pea (Sujkowska *et al*., 2007) and positively regulates nodulation in soybean (Li *et al*., 2015).

The delivery of NPL, SyPME and different cell wall components such as pectins to the IC, the IT tip and the transcellular passage site (Fig. 3 and Fig. S3) fully relies on a well-orchestrated targeted secretion and consequently on an intact cytoskeleton. This is exemplified by the function of API, a SCAR2 protein and subunit of the plant SCAR/WAVE complex, that activates the nucleation of actin filaments (Gavrin *et al*., 2020). In the Medicago *api* mutant, the majority of the ITs is blocked in root cortex cells prior to nodule primordium invasion (Teillet *et al*., 2008). As recently demonstrated, actin dynamics and endomembrane trafficking are impaired in the *api* mutant, which leads to changes in CW properties (Gavrin *et al*., 2020). This is most likely due to the loss of focal secretion of NPL and SyPME to the IT tip and of enzymes such as the CYSTATHIONINE-β-SYNTHASE-LIKE1 (CBS1), which is probably involved in maintaining the cell wall of mature ITs (Sinharoy *et al*., 2016).

## STAR Methods

### Plant growth and hairy root transformation

Seeds of Medicago were washed 6 times with sterile tap water after being sterilized for 20 min with pure sulfuric acid (H_2_SO_4_). The seeds were then treated with bleaching solution (12% NaOCl, 0.1% SDS) for 60s and washed again 6 times with sterile tap water. The sterilized seeds were covered with sterile tap water for 2 hours before being transferred to 1% agar plates and stratified at 4°C for 3 days in darkness. After stratification, seeds were kept in darkness at 24°C for 24 hours for germination. The seed coat was removed from germinated seedlings, which were then used for hairy root transformation as previously described (Boisson-Dernier *et al*., 2001). Transformed seedlings were first placed onto solid Fahräeus medium (containing 0.5 mM NH_4_NO_3_) and incubated in darkness (at 22°C) for three days, following 4 days at 22°C in white light with roots kept in the darkness. One week later, seedlings were transferred onto fresh Fahräeus medium (0.5 mM NH_4_NO_3_) for another 10 days. Afterward, the transformed roots were screened and positive plants were transferred to open pots for phenotyping.

### Phenotyping

Both R108 and *sypme* mutant seedlings were directly grown in open pots (mixture of 1:1 quarzsand:vermiculite mixture, 2 plants/pot) after seed germination. Plants were watered with liquid Fahräeus medium (without nitrate, 30 mL/pot) and tap water (30mL/ pot) once a week, individually. The plants were then inoculated with *S. meliloti* (Sm2011; OD600 = 0.003) 7 days after transfer (20 ml/pot). Another 10 days later, plants were harvested for the quantification of infection structures.

### Visualization of IT phenotypes

Roots transformed with the different NPL-RNAi constructs were harvested 10 days post-inoculation, fixed in PBS solution containing 4 % PFA under vacuum for 15 min (twice) and kept at room temperature for 2 hours before being transferred to a ClearSee solution. Roots were kept in ClearSee for 2-3 days before the solution was refreshed and supplied with 0.1% Calcofluor white prior to imaging.

### *In situ* hybridization

A 228 nucleotides fragment of the SyPME CDS was amplified using specific primers (Table S1) from *Medicago truncatula* A17 cDNA with KOD DNA polymerase (Stratagene, San Diego, CA, USA). The amplified fragment was cloned into a blunt cloning vector pEASY-Blunt3, which worked as the template for generating the Digoxigenin-labelled sense or antisense RNA probes using T7 RNA polymerase. Young nodule primordia (10 dpi) and matured nodules (20 dpi) were collected as material for further in situ hybridizations. Material preparation (fixation, dehydration, infiltration, and embedding) was performed as previously described earlier (Jackson, 1992). Paraffin-embedded materials were sectioned at a thickness of 10 μm for hybridization.

### Cloning and DNA constructs

All initial DNA fragments being used in this study were synthesized by Life Technologies, before being assembled in Golden Gate compatible expression vectors (Weber *et al*., 2011). All designed constructs and related primers used in this study are listed in Table S1. Golden Gate L0 modules for the pLeEXT1 (Rival *et al*., 2012) and pAtPEP (Ron *et al*., 2014; Sevin-Pujol *et al*., 2017) promoters were kindly provided by Dr. Tatiana Vernié (University of Toulouse, France). All Medicago gene sequences were retrieved from the Phytozome database with the gene IDs: *SyPME* (*Medtr4g087980*), *NPL* (*Medtr3g086320*).

### GUS staining

GUS staining of roots and nodules was performed as described previously (Su *et al*., 2020). Stained nodules were embedded in Technovit 7100 for microtome sectioning and sections (10 nm) were further stained with 0.05% w/v Toluidine Blue prior to imaging.

### Immunofluorescence

Nodule sections were blocked with 3% BSA for 30 min at room temperature before adding the primary antibody onto the sections (PlantProbes; 1:50 dilution in PBS solution supplemented with 3% BSA). Incubation with the primary antibody was carried out for 1 h at room temperature or overnight at 4°C in the case of 2F4. Samples were then washed at least three times for 5 min with PBS and the secondary antibody (conjugated with AlexaFluor 561, Sigma-Aldrich; 1:500 dilution in PBS supplemented with 3% BSA) was added for an additional 30 min to one hour incubation in darkness. Finally, the samples were washed at least three times with PBS in the dark prior to image acquisition. All the processes were performed on microscopy glass slides in a homemade humidity chamber at room temperature.

### Fluorescence microscopy

Composite Medicago plants for fluorescence imaging were grown in open pots and inoculated with *S. meliloti* as described above. To study primary infection events, roots were harvested at 7 dpi while nodules were obtained at 2 wpi. Nodules were embedded in 7 % low melting agarose before 70 μm thick vibratome sections were obtained for subsequent confocal microscopy using a Leica TCS SP8 with the following settings: Images were acquired with a 20×/0.75 (HC PL APO CS2 IMM CORR) or a 40x/1.0 (HC PL APO CS2) water immersion objective (only for nodule vibratome sections). Genetically encoded fluorophores were excited using a White Light Laser (WLL) with GFP: 488 nm (ex) /500-550 nm (em); mCherry: 561 nm (ex) / 575-630 nm (em); Calcofluor white: 405 nm (ex; UV Laser) / 425-475 nm (em). Images for CLEM-related immunofluorescence were taken with ZEISS ApoTome.2 light microscope with the following detection settings: CFP: BP 480/40 DMR 25; dsRed: BP 629/62. All the image analyses and projections were performed using either the ImageJ/(Fiji) or Imaris software.

### Sample preparation for TEM analysis

Nodules were sectioned in half and immediately fixed in MTSB buffer containing 8% paraformaldehyde (PFA) and 0,25 % glutaraldehyde (GA) at room temperature under vacuum for 15 min and left in the same fixative solution for 2 h at room temperature. After this, they were further fixed with 4% PFA and 0,125 % GA for 3 h at room temperature and with 2% PFA and 0,65 % GA overnight at 4°C. Nodules were then washed with MTSB buffer, and dehydrated in ethanol graded series at progressive low temperature as follows: 30% EtOH at 4°C 15 min, 50%-70%-95%-and 100% EtOH at -20°C for 15 min each, followed by a second incubation in 100% EtOH at -20°C for 30 min. Embedding in Lowicryl HM20 was performed at -20°C gradually increasing the EtOH/Lowicryl ratio (1:1 for 1h, 1:2 for 1h, 0:1 for 1h). After the last step, the resin was replaced with fresh Lowicryl and the samples were incubated overnight at - 20°C. For polymerization, the resin was replaced once more, and the samples were kept at - 20°C for 2 days under UV light. Blocks were sectioned with a Reichert-Jung Ultracut-E microtome. Ultrathin (70 nm) sections were collected in copper slot or finder grids and observed in a Philips CM10 (80 kV) microscope coupled to a GATAN Bioscan Camera Model 792 or a Hitachi 7800 TEM coupled to a Xarosa CMOS camera (Emsis).

### Sample preparation for CLEM

CLEM was applied on 70 nm Lowicryl HM20 ultrathin sections obtained with a Reichert-Jung ultramicrotome and collected on finder grids. The grids with the sections were washed with PBS buffer for 5 min, followed by an incubation with 0.12 M Glycine in PBS for 10 min. After 5 min washing in PBS, the grids were incubated for 10 min in blocking solution (3% BSA in PBS) followed by 30 min incubation with the first antibody in blocking solution. After six times washing for 3 min each in PBS, the grids were incubated for 30 min in blocking solution containing the above-mentioned fluorescence labelled secondary antibody. Grids were washed six times for 3 min each in PBS and incubated in 1% DAPI solution for 5 min before they were mounted on a microscope glass slide for observation with a fluorescence microscope.

### Immuno-Gold staining

Samples for immuno-gold staining were treated as for CLEM but substituting the secondary antibody by conjugated Protein A-gold 5 nm (University Medical Center Utrecht) or Goat-anti Rat IgG 12 nm gold conjugated polyclonal antibody (Jackson Immuno Research (112-205-167)), and contrasting the sections with 2% uranyl acetate after washing in water.

## Acknowledgements

We would like to thank Eija Schulze, Rosula Hinnenberg and Carmen Schubert for their experimental support and technical help and the entire Ott lab team and Elke Barbez for the continuous discussions on the project. We also thank the staff of the Life Imaging Center (LIC) in the Hilde Mangold House (HMH) of the Albert-Ludwigs-University of Freiburg for help with their confocal microscopy resources, and the excellent support in image recording. The microscopes are operated by the Microscopy and Image Analysis Platform (MIAP) and the Life Imaging Center (LIC), Freiburg. Norbert Roos and Jens Wohlmann (Electron Microscopy Facility, Department of Biosciences, University of Oslo, Norway) are acknowledged for helping with the SEM sample preparation and imaging. The *Medicago truncatula* plants utilized in this research project, which are jointly owned by the Centre National De La Recherche Scientifique, were obtained from Noble Research Institute, LLC and were created through research funded, in part, by a grant from the National Science Foundation, NSF-0703285.

## Authors contributions

Conceptualization: CS, XL, TO; Investigation: CS, GZ, MR-F, JW, WY, PL, LU; Writing – Original Draft: CS, TO; Writing –Review & Editing: CS, GZ, MR-F, JW, WY, PL, LU, XL, TO; Supervision: CS, XL, TO; Project administration: XL, TO; Funding Acquisition: CS, GZ, XL, TO

