## Supplemental Figures for "Transcellular progression of infection threads in *Medicago truncatula* roots is controlled by locally confined cell wall modifications"

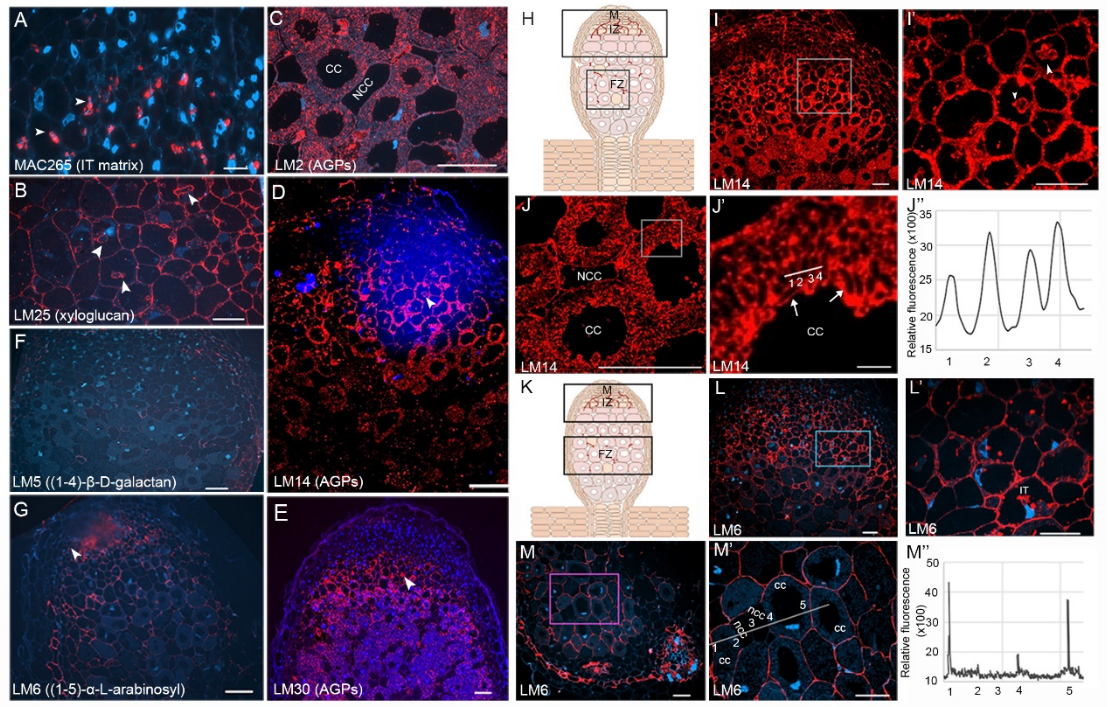


**Fig. S1: Immunofluorescence labeling with different cell wall antibodies.** (A-G) 14 days old Medicago nodules were embedded in Lowicryl HM20 and hybridized with the respective antibodies (red). Cellular structures were labelled by different antibodies: (A) IT matrix (MAC265); (B) xyloglucan (LM25); arabinogalactan proteins (AGPs; LM2 (C), LM14 (D), LM30 (E)); different types of RG-I (LM5 (F) and LM6 (G)). DNA was counterstained with DAPI. Arrowheads indicate infection threads. Scale bars (A-G) indicate 50 µm. (H-J'')14 days old Medicago nodules were embedded in Lowicryl HM20 and hybridized with the LM14 antibody (red). (H) Sketch of a Medicago nodule with boxes indicating regions shown in the respective panels. (I-J') Robust labelling of the infection zone (I-I') and infected cells of the fixation zone (J-J'). Boxes (in I and J) indicate the close-ups shown in (I') with ITs marked by an arrowhead and of infected cells (J') with symbiosomes marked by arrows. Line (in J') indicates area of a transect to illustrate fluorescence intensities (J''). Scale bars indicate 50 µm in (I-J), and 5 µm in J'. (K-M'') 14 days old Medicago nodules were embedded in Lowicryl HM20 and hybridized with the LM6 antibody (red). (K) Sketch of a Medicago nodule with boxes indicating regions shown in the respective panels. (L-M') Concise labelling of all nodular cell types of the infection (L-L') and fixation zone (M-M'). Boxes (in L and M) indicate the close-ups shown in (L') with additional labelling of infection threads (IT) and of infected cells where only the peripheral but not the symbiosome membrane was labelled (M'). Line in (M') indicates area of a transect to illustrate fluorescence intensities (M''). Scale bars (L-M') indicate 20 µm. M: meristem; IZ: infection zone; FZ: fixation zone; CC: colonized cell; NCC: non-colonized cell.

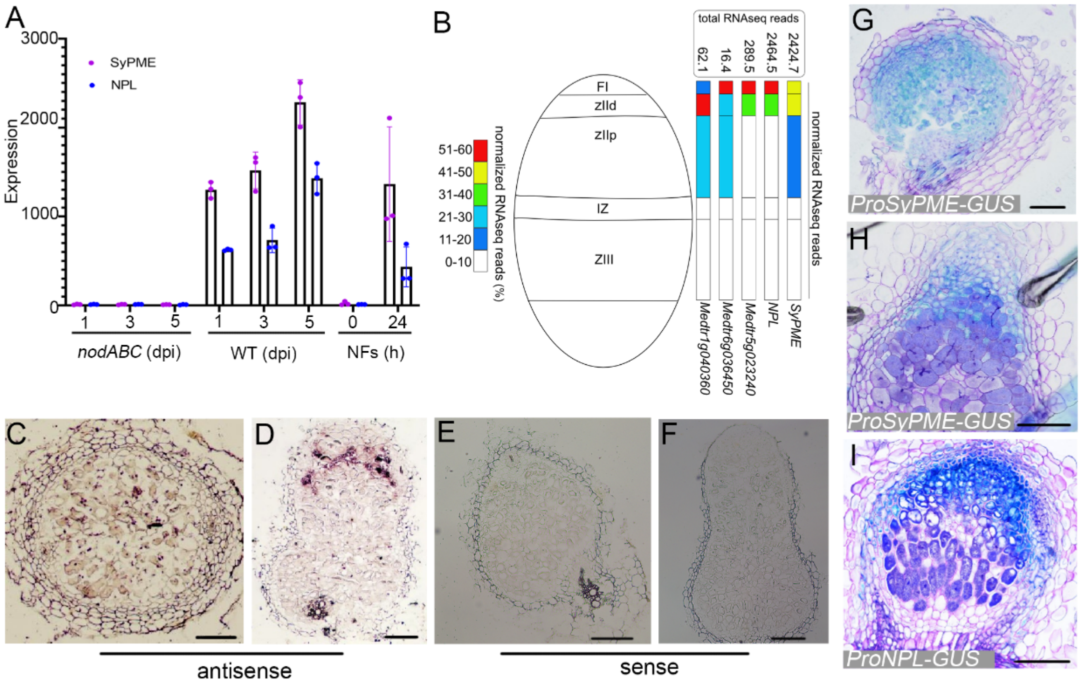


**Fig. S2: Analysis of *SyPME* expression domains.** (A) Expression patterns of NPL and SyPME are highly correlated (0.9883) as exemplified upon inoculation of roots with a Nod Factor (NF)-deficient *S. meliloti nodABC* strain (*nodABC*), WT *S. meliloti* and isolated NFs at different time points. Data were retrieved from Breakspear et al., 2014. Dpi: days post inoculation; h: hours. (B) Schematic overview of the spatial expression of SyPME and different pectate lyases including NPL in different zones of an indeterminate Medicago nodule. Original data were retrieved from Roux et al., 2014. FI: nodule meristematic zone; zIId: distal of infection zone; zIIp: proximal of infection zone; IZ: interzone; ZIII: nitrogen-fixation zone. (C-F) Spatial analysis of *SyPME* transcript accumulations by i*n situ* hybridization using a *SyPME* antisense (C-D) and sense (control) probe (E-F) on 14 days old Medicago nodules. Magenta precipitates indicate presence of *SyPME* mRNA. (F-H) Promoter GUS (blue) analysis for *ProSyPME* (G-H) and *ProNPL* (I) on 14 days old transformed *M. truncatula* nodules counterstained with Toluidine Blue (purple). Scale bars indicate 50 µm.

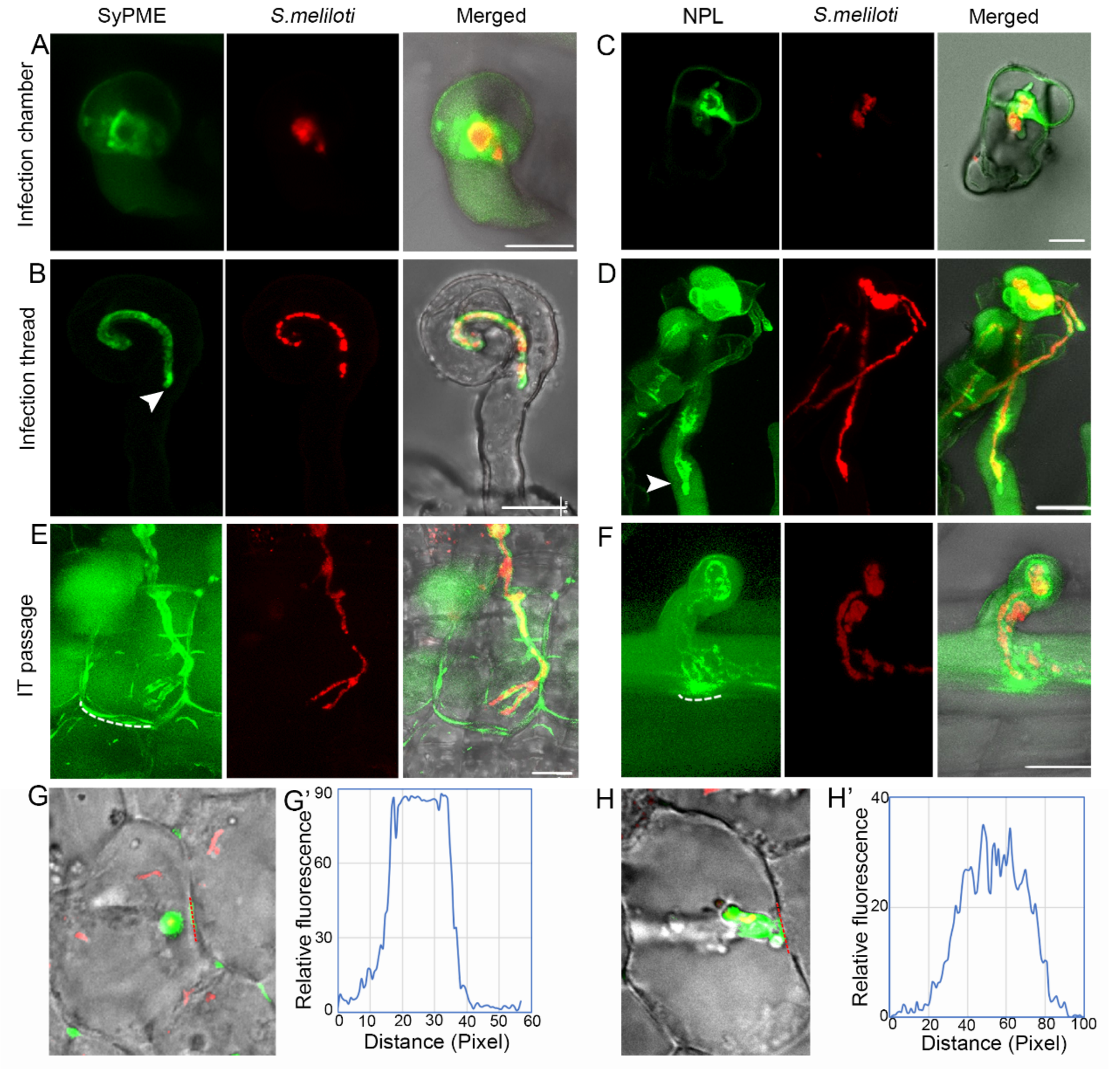


**Fig. S3: SyPME and NPL accumulate in spatially overlapping sites.** (A-B) A SyPME-GFP fusion protein (green) localizes to the infection chamber (IC; A) and along growing infection threads (IT) including the IT tip (arrowhead) (B). (C-D) Similar patterns were observed for NPL-GFP (green) at the IC (C) while NPL predominantly localized to the IT tip (arrowhead) and young parts of the IT (D). Initial labelling of future transcellular passage sites prior to IT passage was found for SyPME (E) and NPL (F) as indicated by the dashed line. Fluorescent *S. meliloti* is shown in red. All images are maximal projections. Scale bars indicate 10 µm. (G-H') SyPME and NPL are accumulate at future passage sites prior to IT arrival. SyPME (G) and NPL (H) accumulate at prospective passage sites and at the tip of IT (images are identical with panels Fig. 3E and Fig. 3H). Dashed red line indicates transect used to determine fluorescence intensities at these sites for SyPME (G') and NPL (H').

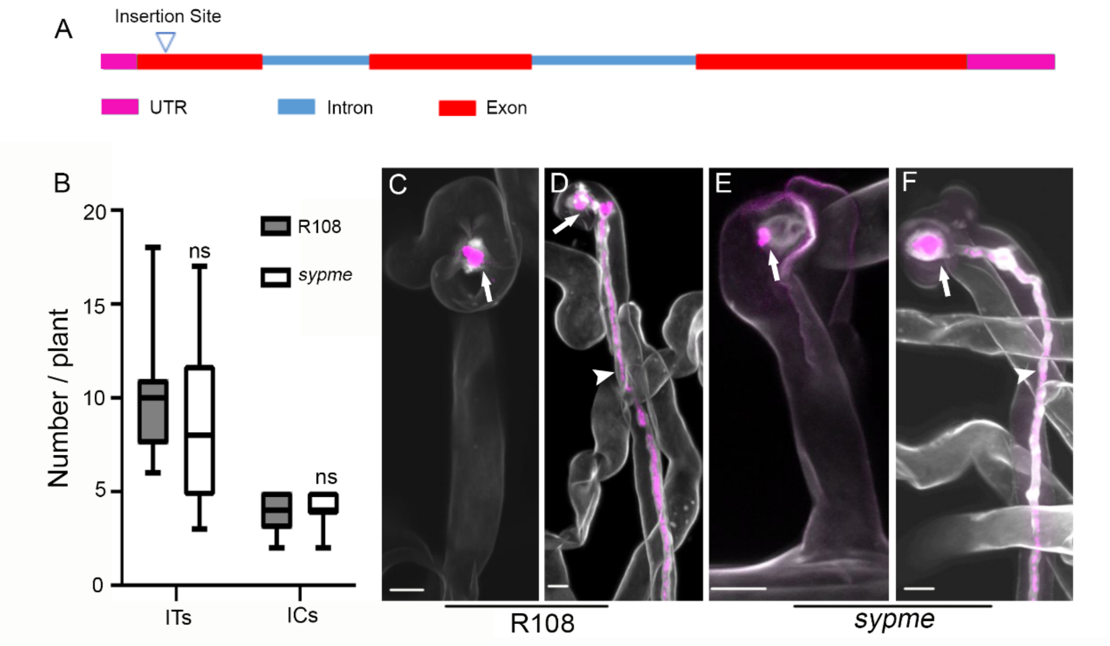


**Fig. S4: A single locus *sypme* mutant does not exhibit a symbiotic phenotype.** (A) Schematic representation of the *SyPME* gene structure and mapped Tnt1 transposon insertion site (line NF2281). UTR, untranslated region. (B) Infection chambers (ICs) and infection threads (ITs) were scored at 10 days after inoculation, with *n* =10 root systems for each genotype. Data are means ± SE. Statistics were performed using an unpaired two-tailed t-test. ns: Not significant. (C-F) IC and IT morphology were visualized by Calcofluor-white staining (White color) in R108 (C-D) and *sypme* mutants (E-F). Fluorescent *S. meliloti* is shown in magenta. Scale bars indicate 10 µm. Images (C-F) are maximal projections.
