## Supplemental Table for "Transcellular progression of infection threads in *Medicago truncatula* roots is controlled by locally confined cell wall modifications"

**Table S1: List of antibodies and their putative target epitopes used in this study**

| Antibody name | Epitope |
|---|---|
| MAC265 | Infection thread matrix glycoprotein |
| LM5 | Pectic polysaccharide ((1-4)-β-D-galactan) |
| LM6 | Pectic polysaccharide ((1-5)-α-L-arabinosyl) |
| LM19 | Un-esterified homogalacturonan |
| LM20 | Methyl-esterified homogalacturonan |
| 2F4 | 'egg box" dimer conformation of homogalacturonan |
| LM2 | Arabinogalactan-proteins (AGPs) |
| LM14 | Arabinogalactan-proteins (AGPs) |
| LM30 | Arabinogalactan-proteins (AGPs) |
| LM25 | xyloglucan |
